# HT SpaceM: A High-Throughput and Reproducible Method for Small-Molecule Single-Cell Metabolomics

**DOI:** 10.1101/2024.10.24.620114

**Authors:** Jeany Delafiori, Mohammed Shahraz, Andreas Eisenbarth, Volker Hilsenstein, Bernhard Drotleff, Alberto Bailoni, Bishoy Wadie, Måns Ekelöf, Alexander Mattausch, Theodore Alexandrov

## Abstract

Single-cell metabolomics promises to resolve metabolic cellular heterogeneity, yet current methods struggle with detecting small molecules, throughput, and reproducibility. Addressing these gaps, we developed HT SpaceM, a high-throughput single-cell metabolomics method with novel cell preparation, custom glass slides, small-molecule MALDI imaging mass spectrometry protocol, and batch processing. We propose a unified framework covering essential data analysis steps including quality control, characterization, differential analysis, structural validation and functional analysis. Interrogating human HeLa and mouse NIH3T3 cells, we detected 73 diverse small-molecule metabolites validated by bulk LC-MS/MS, achieving high reproducibility across wells and slides. Interrogating nine NCI-60 cancer cells and HeLa, we identified cell-type markers in small subpopulations. Functional analysis revealed overrepresented metabolic pathways, co-abundant metabolites, and metabolic hubs. We demonstrate the ability of SCM to analyze over 120,000 cells from over 112 samples, and provide guidance to interpret single-cell metabolic heterogeneity, revealing metabolic insights beyond population averages.

## Introduction

Single-cell omics are revolutionizing biology by shedding light on cellular heterogeneity, by revealing novel cell types, functional phenotypes, and states, as well as variability of cellular programs within these cell subsets ^1^. Detecting and understanding this heterogeneity is challenging, yet paramount to understand homeostasis, disease onset, progression, responses to therapies and environmental stimuli ^2,3^. Single-cell metabolomics (SCM) rapidly emerged as a technology of choice to probe the metabolic underpinnings of this heterogeneity by directly detecting the end-products of cellular metabolism ^4–7^. In addition to fluorescence-based assays, Raman spectroscopy, and nuclear magnetic resonance, mass spectrometry (MS) has emerged as the major approach for SCM, due to its sensitivity and specificity ^8^. Among a variety of mass spectrometry methods, using Matrix-Assisted Laser Desorption Ionization (MALDI)-imaging mass spectrometry, a technology originally developed for spatial metabolomics ^9,10^, is becoming increasingly widespread for SCM due to its commercial availability, rapid technological progress, speed, sensitivity, and potential to extrapolate analyses to tissues sections ^6^.

However, current MALDI-imaging-based SCM methods struggle with detecting small-molecule metabolites with most reports focusing on detecting highly-abundant and easily-ionizable lipids, in particular phospholipids composing the cell membrane. Saunders et al. ^11^ in Table 1 alludes to SCM laser-based imaging mass spectrometry only reporting lipids as metabolites. Confirming this review, our search through original papers and preprints from 2024 with the keywords “MALDI” and “single-cell metabolomics” (as of October 2024) showed that five ^12–16^ out of six publications reported lipids only, where the only one reporting small-molecule metabolites ^17^ used a custom MALDI-2 postionization setup. This reflects the challenges in detecting small-molecule metabolites in single cells due to their low abundance yet lack of amplification, high structural diversity, wide dynamic range, and propensity to cellular leakage during sample preparation. However, a robust detection of small-molecule metabolites is needed for a majority of metabolism studies, and the lack of it limits the applicability of SCM and impedes its impact in biology and pharmacology.

The second challenge hindering the uptake of SCM is the relatively low throughput in terms of the number of samples and cells, and a low reproducibility. This is particularly important because SCM has a strong potential for high-throughput due to its relatively low per-sample cost. as the reagents are substantially cheaper compared to sequencing-, probe- or antibody-based single-cell omics. This need is actively discussed with multiple advances achieved recently ^18^. Despite MALDI-imaging providing the highest detection rate in SCM compared to other approaches (see Table 1 in ^18^), the numbers of reported cells remain relatively low, often below 10,000 cells. Cell throughput is often limited by the variability in sample preparation, batch effects preventing integrating data across multiple slides, and a high prevalence of sample-outliers and cell-outliers. Among the largest studies in SCM, we found reports of profiling 30,000 ^19^ and 30,584 ^20^ single cells, both using MALDI-imaging. Yet, there is a need for robust high-throughput SCM methods for interrogating larger numbers of samples and cells, first for increasing the confidence in SCM by routinely using more replicates and, second, for applying SCM in larger biological and clinical studies.

The third challenge in SCM is the lack of established frameworks and guidelines for data analysis. In particular, there is not clear guidance for quality control, assessing variability among replicates, or quantifying variations in metabolite levels. This is a frontier of SCM reflected by only a few SCM-focused reviews highlighting it as an important future step towards the maturity of the field ^6,7^. Yet, there is an increasing appreciation of the need for establishing data analysis frameworks ^21,22^.

In this work, we aim to address these challenges by proposing a new SCM method, HT SpaceM, and a computational framework and guidelines for SCM data analysis. HT SpaceM follows the principles of the recently published SCM method SpaceM ^20^ yet with new cell preparation focusing on small-molecule metabolites, custom laser-etched glass slides enhancing microscopy and image analysis, new MALDI-imaging protocol optimized for detecting small-molecule metabolites, and batch processing. The data analysis framework includes quality control and assessment of data variability, single-cell characterization, structural validation, differential analysis, and network analysis. We validated the method and its reproducibility by analyzing HeLa and NIH3T3 cell lines on 3 glass slides, detected 135 ions in 78,500 cells across 72 samples, with 73 metabolites validated by Liquid Chromatography Tandem MS (LC-MS/MS) bulk metabolomics. By analyzing a subset of nine NCI-60 cancer cell lines and HeLa cells (202 ions in 42,153 cells across 40 samples), we identified cell-line-specific metabolic markers, discovered co-abundant metabolites and metabolic hubs through single-cell co-abundance and network analysis.

## Results

### HT SpaceM method

We present HT SpaceM, a method for high-throughput small-molecule single-cell metabolomics using MALDI-imaging mass spectrometry. HT SpaceM can analyze up to 40 samples (different cell types or replicates of those) plated on the same glass slide, detects over 1,000 cells from each sample and is applicable to cells of different types. HT SpaceM follows the principles of SpaceM ^20^, a method we published previously, and uses MALDI-imaging similar to microMS ^23^ and MAMS ^24^.

The experimental steps of HT SpaceM start with laser-etching the glass slides with well-identifiers and fiducials (**Figure 1A**). The cells are plated into removable chambered wells mounted onto a glass slide with 40 out of 64 wells available for analysis as being within the area of acquisition of the AP-SMALDI5-imaging system used (**Figure 1B**). After adherence, cells can be fixed or fluorescently stained for improved cell segmentation or phenotyping. After removal of the chambers, washing is performed to remove the cell culture medium, followed by the cells desiccation under vacuum. The first round of microscopy (brightfield and fluorescent) delivers cell outlines and phenotypes (**Figure 1C**). MALDI-imaging mass spectrometry is performed with a protocol enhancing detection of small-molecule metabolites. Finally, the second round of microscopy (brightfield) delivers MALDI laser ablation marks. Following registration of pre- and post-MALDI microscopy images, overlay of cell outlines and ablation marks positions is performed.

**Figure 1.**
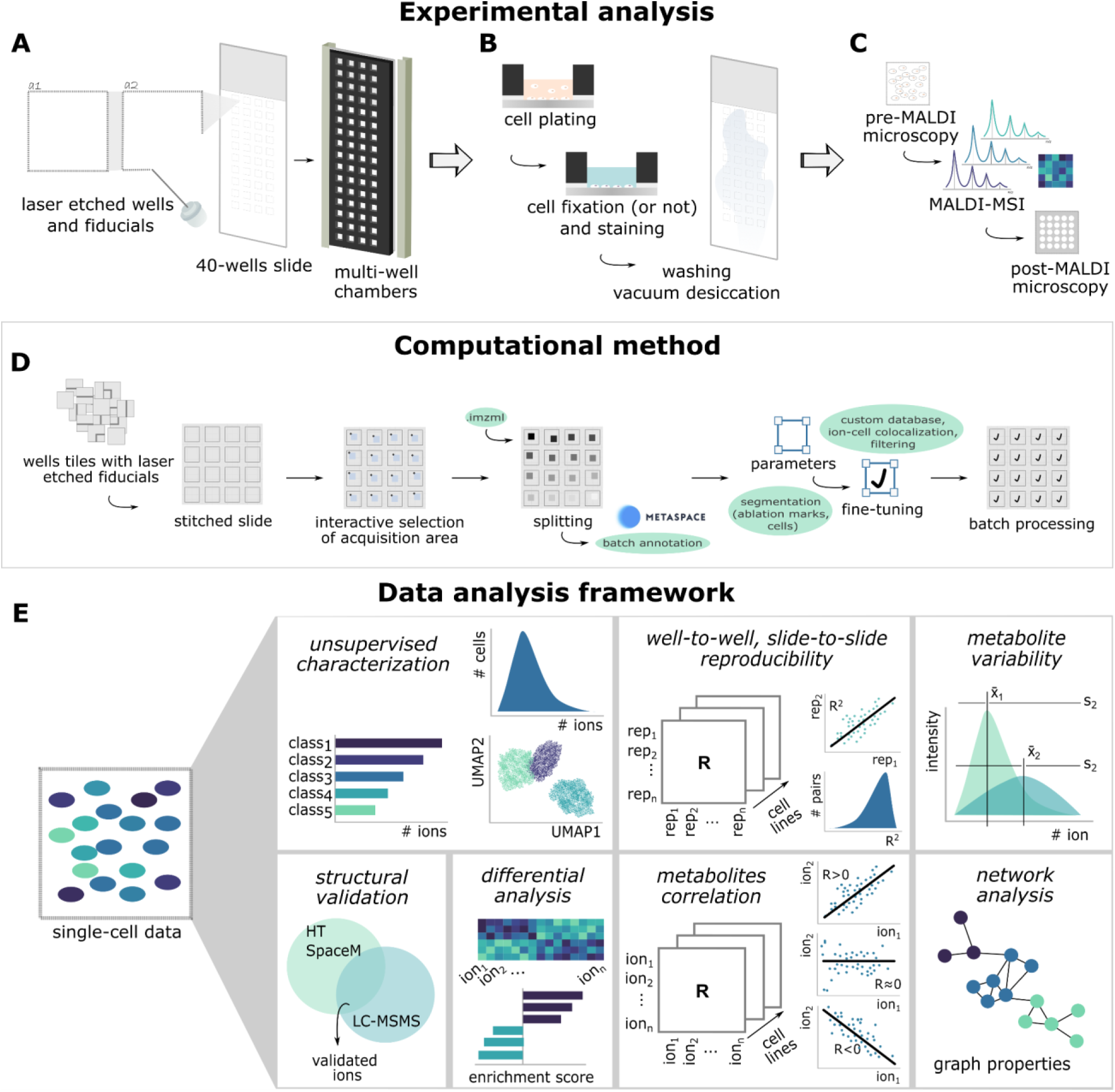
The High-Throughput SpaceM method workflow and the proposed single-cell data analysis framework. (**A**) Fiducial marks and well-identifiers are laser-etched onto a glass slide, with removable multi-well chambers added. (**B**) Cells are plated into wells to allow cell adhesion followed by the optional addition of fluorophores for cell staining; cells can be further fixed prior to washing with volatile solution. (**C**) The entire slide is imaged using brightfield and fluorescence microscopy before application of a MALDI matrix. MALDI-imaging MS is performed followed by a new round of brightfield microscopy (post-MALDI) to visualize the laser ablation marks. (**D**) Fiducials facilitate stitching overlapping microscopy image tiles, registration for pre- and post-MALDI microscopy images, and facilitating interactive selection of MALDI-imaging acquisition areas within every well. MALDI-imaging files are batch-uploaded to METASPACE for metabolite annotation. Batch pixel-cell deconvolution is performed using fine-tuned parameters. (**E**) Single-cell data analysis framework for SCM comprising unsupervised characterization of conditions, assessment of reproducibility among wells and slides, and metabolite variability, structural validation of annotations with bulk LC-MS/MS, identification of differentially expressed markers, and single-cell metabolites co-abundance analysis through correlation and networks.

The computational steps of HT SpaceM (**Figure 1D**) start with stitching the microscopy tiles to obtain a wide-field image of the whole slide. Using in-house software, for each well we select the best area for MALDI-imaging presenting the best cell confluency and lack of visible washing artifacts. Following automated MALDI-imaging across all wells, the resulting raw file is centroided and split into individual imzML files, one for each well. The imzML files are submitted to METASPACE ^25^ for batch metabolite annotation against the HMDB v4 ^26^ and CoreMetabolome v3 databases ^27^. After pulling resulting annotations, we create a custom metabolite database comprising ions co-localized with cells. Each well-dataset is reannotated on METASPACE against this custom database to reduce the dataset-wide dropouts. In parallel, cells in the pre-MALDI microscopy images are segmented using the Cellpose *cyto2* model ^28^ fine-tuned on individual cells from the experiment (**Figure 2A**). Pixel-cell deconvolution, normalization, finding intracellular ions, and cells and ions filtering is performed as in SpaceM ^20^, delivering single-cell metabolic profiles.

**Figure 2.**
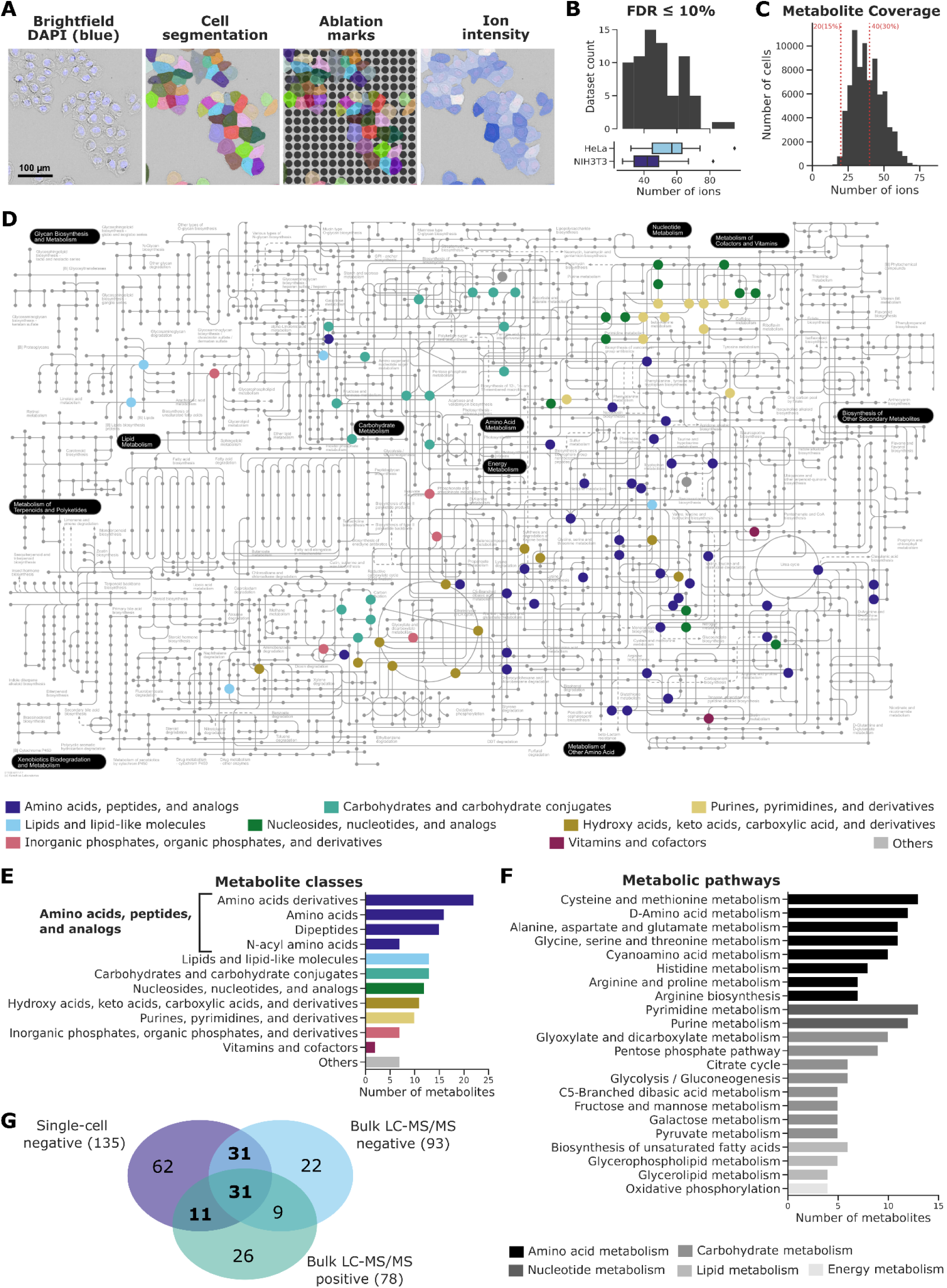
Comprehensive coverage of major metabolic pathways and small-molecule metabolite classes by HT SpaceM. (**A**) Brightfield-DAPI microscopy image of HeLa cells highlighting cell segmentation using Cellpose, the overlay of cells and laser ablation marks, and single-cell intensities of the ion of N-acetylaspartate [C_6_H_9_NO_5_-H]^−^ after HT SpaceM processing. (**B**) Number of ions [M-H]^−^ per well at FDR≤10%. (**C**) Metabolite coverage of single-cell profiles of HeLa (66.2% cells) and NIH3T3 (33.8% cells). (**D**) The KEGG map of primary human metabolism highlighting detected endogenous metabolites (nodes) colored by molecular class, generated in iPath3 ^33^. Lines represent metabolic reactions. (**E**) Barplot showing the detected metabolite classes. (**F**) Barplot showing the detected human metabolic pathways. (**G**) Venn diagram highlighting the total number of molecular formulas detected by HT SpaceM (negative ion mode) and bulk LC-MS/MS (negative and positive ion modes).

Using new multi-well chambers in HT SpaceM increases the throughput five-fold in the number of samples compared to SpaceM. By using smaller chambers for cell plating, HT SpaceM method enhances reproducibility and reduces batch effects, as more samples per condition can be analyzed in a single experimental run. Moreover, smaller well sizes require fewer seeded cells, better aligning with the effective MALDI acquisition area, and increasing cell management efficiency. Using laser-etched slides allowed us to substantially simplify and streamline microscopy tiles stitching, facilitating the registration of microscopy images, and to use the whole-slide overview image for selecting the best areas for MALDI analysis. Using Cellpose with fine-tuning allowed us to reduce the cell segmentation time considerably. Compared to SpaceM, we use a grid of spots (here called grid fitting) to manually find the exact positions of laser ablation marks in each MALDI area. Grid fitting showed to be much faster, more robust, better suited for small MALDI step sizes which often results in overlapping ablation marks. This improvement was made possible with a high-accuracy machine stage in AP-SMALDI5, compared to the AP-SMALDI10 system used in SpaceM.

Data handling was streamlined by using the SpatialData framework ^29^ and increased use of METASPACE for automated metabolite annotation and reannotation, as well as for quality control, data storage, sharing, and publication.

### Data analysis framework

We present and showcase a data analysis framework covering steps we found to be essential for analysis of SCM data: quality control, unsupervised characterization, evaluation of reproducibility, quantification of metabolite variability, and structural validation of metabolites annotated in MALDI-imaging data (**Figure 1E**). Moreover, we present a novel approach to interpretation of SCM data with a functional analysis of markers covering overrepresented metabolic classes and pathways, and a single-cell metabolic network analysis based on calculating metabolite co-abundance across single cells.

### HT SpaceM achieves comprehensive small-molecule metabolites coverage across diverse metabolic classes and pathways

Despite recent advances in mass spectrometry, detection of small-molecule metabolites in SCM is still challenging, due to loss of intracellular metabolites during sample preparation, low sample volume and limited sensitivity, and ion suppression favoring detection of metabolites of highest ionization efficiency ^11^. Many SCM methods, including our original SpaceM publication, report mainly lipid detection over small molecules due to lipids’ higher abundance and easier detectability. This is common not only for MALDI-imaging-based SCM, since in ESI-based SCM, reported ranges of 5-500 metabolites often include lipids as well ^7,8^.

To assess HT SpaceM small-molecule metabolites coverage, we utilized HeLa and NIH3T3 cell lines as models of different mammalian cells with distinct morphological phenotypes and metabolic profiles. We considered three slides, one slide where MALDI-imaging was acquired for all well-replicates of HeLa and NIH3T3, and two other slide-replicates partially acquired, with 72 wells analyzed in total. Consolidating metabolite annotations from MALDI-imaging datasets, we obtained 135 small-metabolite ions. This number is higher than typically found in spatial metabolomics studies where, according to the METASPACE knowledge base ^30^, an average of 26 metabolites or lipids are detected (median, at FDR10%). In either HeLa or NIH3T3 cells, we annotated 30-100 ions per well at FDR 10% (**Figure 2B**). HeLa cells consistently exhibited more detected ions than NIH3T3 cells for all considered FDR levels (**Suppl. Figure 1B**). In single-cell data, we detected 20-80 ions per cell after deconvolution and metabolite curation (**Figure 2C**). Notably, we observed cell heterogeneity in the numbers of detected ions, with only 10% of ions present in more than 90% of cells, and 47% of ions present in less than 10% of the cell population. The experiment resulted in an overall detection of 135 [M-H]^−^ ions, corresponding to endogenous metabolites co-localized with cells.

The detected ions play crucial roles in human metabolism. For the detected 135 ions, we found 101 metabolites from human metabolic pathways described in the KEGG database (**Figure 2D**). This coverage extends to diverse metabolite classes (**Figure 2E**), with amino acids, peptides, and analogs (n=60) being the most frequent class. The lipid class mainly comprises fatty acids and small lipid derivatives due to the set *m/z* range of 100-400 and their ability to get ionized in negative mode. Importantly, the represented metabolite classes align with the expected metabolite detectability in experiments using AP-MALDI-imaging MS ^31^. Multiple metabolic pathways are represented, with the most metabolites from cysteine and methionine, pyrimidine, purine, and amino acid metabolism (**Figure 2F**).

The limited metabolite abundance in single cells poses challenges for conducting untargeted MS/MS experiments *in situ*. We opted to employ bulk LC-MS/MS of cell lysates to refine and validate the metabolites putatively identified in the samples based on MALDI-imaging MS1 as commonly done in SCM. The bulk LC-MS/MS data revealed 151 metabolites, corresponding to 130 molecular formulas (**Figure 2G**). A substantial overlap between HT SpaceM coverage and bulk LC-MS/MS data was observed, with around 54% (n=73) of the molecular formulas from SCM detected by both techniques. LC-MS/MS was conducted in negative and positive ionization modes, and validation was performed by matching formulas detected in SCM data to metabolites identified in the bulk LC-MS/MS, resulting in 74 identified metabolites at the Level 1 and 17 putatively annotated metabolites at the Level 2 as recommended by the Metabolomics Standards Initiative ^32^. These shared metabolites span seven of eight main molecular classes, consistently detected and validated by both bulk and single-cell approaches (**Suppl. Figure 1C**).

Evaluating the throughput, we obtained metabolic profiles for 78,500 single HeLa or NIH3T3 cells that represent 1,090 cells per well, on average. The total number of cells detected varied depending on cell type, size, and morphology, affecting confluency and cell spread after seeding. For example, HeLa cells, being smaller than NIH3T3 fibroblasts (**Suppl. Figure 1A**) and growing in clusters, resulted in more HeLa cells detected (52,006) than NIH3T3 cells (26,494). The demonstrated performance of HT SpaceM substantially exceeds the capacities of SpaceM, which detects mainly lipids, in handling five times more samples per slide. Among comparable methods, microMS covered mainly lipids and neurotransmitters, with the largest number of cells reported being 30,000 ^19^, and MAMS, which reports the numbers of cells of 2 to 3 orders of magnitude lower than HT SpaceM.

Overall, HT SpaceM addresses a gap in SCM by detecting small-molecule metabolites encompassing major metabolite classes and pathways, including amino acids, nucleotides, fatty acids, and carbohydrates^20^. Importantly, the majority of metabolites annotated in HT SpaceM were validated by bulk LC-MS/MS, which detected metabolites of similar molecular diversity.

### HT SpaceM demonstrates high reproducibility across wells and slides with low variability for most of detected metabolites

Achieving high reproducibility alongside high sensitivity is essential for emerging technologies like SCM. However, assessing reproducibility is challenging when constrained by the number of cells and technical replicates. Moreover, the lack of established approaches for quantifying reproducibility in SCM represents a significant challenge, as discussed in ^6,21,34^. Here, we propose a data analysis framework to quantify reproducibility across well-replicates and slide-replicates and to calculate the variability of individual metabolites.

We computed mean ion intensity for each metabolite in a well-replicate and calculated Pearson correlation between well-replicates to evaluate well-to-well reproducibility (**Figure 3A**). Intra-slide replicates exhibited high reproducibility for each cell line (**Suppl. Figure 2A**), indicating consistent linear correlation across replicates. Approximately 76% of intra-slide well-well pairs exhibited a coefficient of correlation (R)≥0.97; the lowest values were observed for HeLa cells in slide 2. As expected, inter-slide replicate reproducibility was lower than intra-slide results, yet still with high R values (**Figure 3B**). Inter-slide Pearson R ranged from 0.88 to 0.99, with 61% of the paired replicates having R≥0.97. Well-to-well reproducibility was consistent across intra- and inter-slide replicates (**Figure 3C**), irrespective of the cell line. Slopes of the best fitting line between wells added another layer for reproducibility assessment, where a small deviation from 1 suggests the overall intensities achieved in each well were similar. Around 63% of the paired replicates had slopes≥0.97; 58% for interslide replicates and 70% for intra-slide replicates (**Suppl. Figure 2B**). Zooming in onto well-to-well reproducibility, no replicate outliers were identified in slide 2 (**Suppl. Figure 2C**), with replicates presenting overall similar ion intensities, leading to the clustering of replicates within the cell line. Moreover, no batch effects were observed for either the well position (rows A-J; columns 1-4) or MALDI acquisition order. We also calculated mean ion intensities per slide and cell line, achieving high slide-to-slide reproducibility, with Pearson Rs exceeding 0.98 (**Figure 3D**).

**Figure 3.**
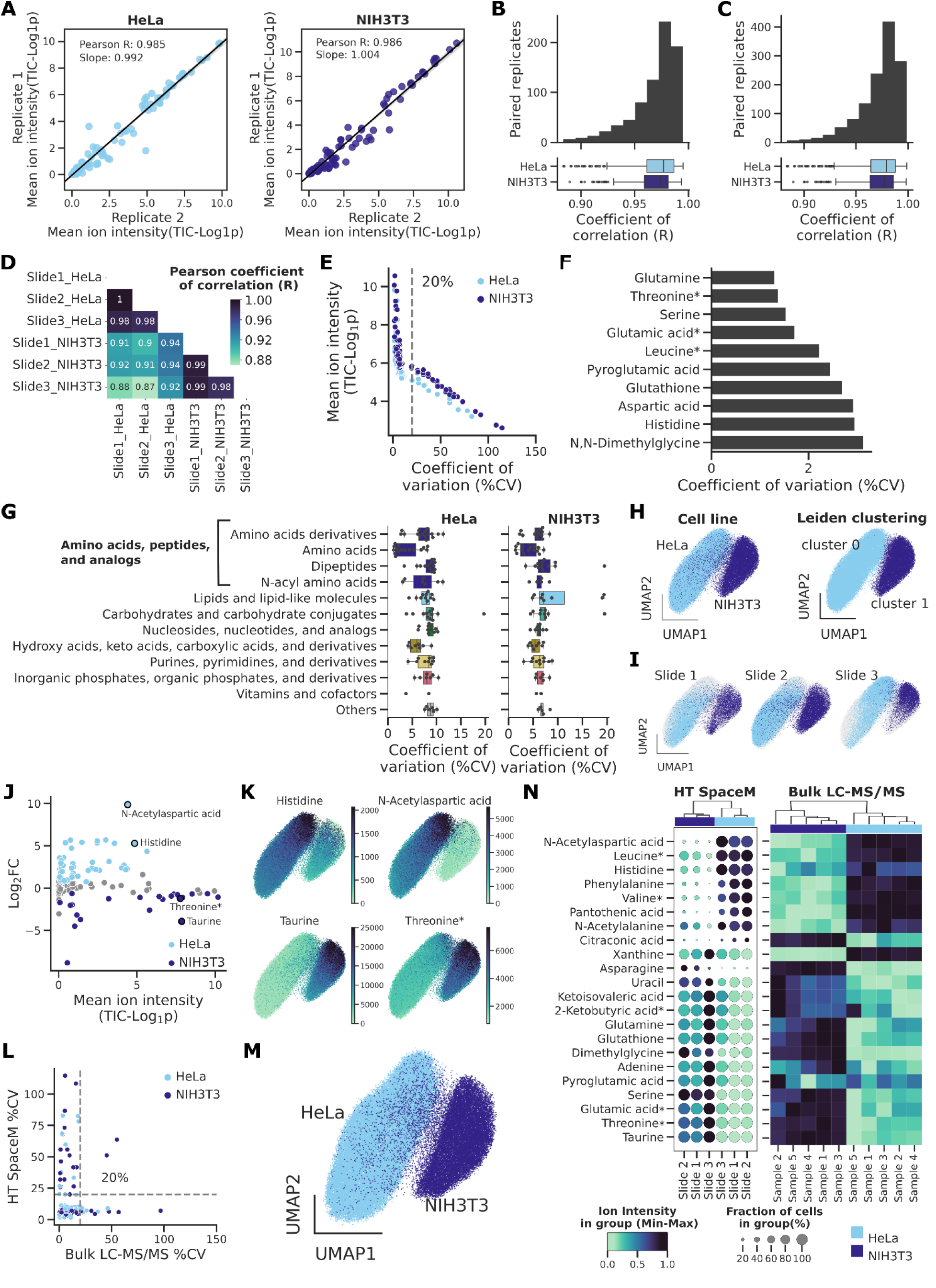
HT SpaceM demonstrates high reproducibility across wells (n=72 wells) and low metabolite variability, delivering metabolite detection comparable to bulk LC-MS/MS. (**A**) Well-to-well reproducibility demonstrated by Pearson Coefficient of Correlation (R) and slope, calculated for the ion intensities observed between intra-slide well-replicates, separately for HeLa and NIH3T3. (**B**) Distribution of Pearson R calculated between replicates within the cell line (HeLa, NIH3T3) from different slides. (**C**) The same as in (**B**) but for all well-replicates (both inter- and intra-slide). (**D**) Pearson R showing slide-to-slide reproducibility. (**E**) Mean ion intensity within condition (non-zero cell fraction) plotted versus the metabolite Coefficient of Variation (%CV) for all detected single-cell metabolites. (**F**) Metabolites with the smallest %CV (mean across HeLa and NIH3T3) among all single-cell detected ions. (**G**) Metabolites %CV in HeLa and NIH3T3 for different molecular classes, considering all formulas detected in single-cell data. (**H**) UMAP of the single-cell data from HeLa and NIH3T3 cells considering all detected metabolites (n=135); figure colored by cell line (left) or Leiden clustering (right). (**I**) The same as in (**H**) but showing data from individual slide-replicates (**J**) MA plot for the distribution of fold change (Log_2_FC) and average ion intensity between HeLa and NIH3T3. Differential analysis resulted in significant metabolites (*p*-value<0.05, FDR-adjusted) with FC≥1.5 highlighted in light blue if increased in HeLa and dark blue if increased in NIH3T3. (**K**) UMAP colored by normalized intensities of differentially expressed markers. (**L**) %CV of single-cell (non-zero cells fraction) and bulk analysis for common ions validated at Level 1 or 2. (**M**) UMAP of the single-cell data but only for ions validated by bulk LC-MS/MS (Level 1 or 2) and with %CV≤20% (n=59). (**N**) HT SpaceM (dot plot) and bulk LC-MS/MS data (heat map) for a subset of differentially expressed metabolites. Dot plot grouped by cell line and slide showing the fraction of non-zero ion intensity cells and feature-scaled mean ion intensity. Heat map showing feature-scaled average metabolite abundance per cell line and replicate. All displayed molecule names were validated in bulk LC-MS/MS at the Level 1, with (*) indicating when isomers were present.

We further examined variability of intensities for individual metabolites across replicates by calculating the Coefficient of Variation (%CV). Our analysis revealed a wide range of ion %CVs (**Figure 3E**) spanning from as low as 1.2% (glutamine) to as high as 82.8% (creatinine) for HeLa cells, while the same metabolites displayed different %CVs in NIH3T3 cells, 1.4% and 32.0%, respectively. We noted that high-intensity ions generally showed less variability among replicates (**Figure 3E**), with the Total Ion Count (TIC) normalization assisting in decreasing ion variability (**Suppl. Figure 2D**). Overall, most ions demonstrated a low %CV. Specifically, 81.48% of the detected ions had %CVs≤20%, corresponding to 88.17% of HeLa ions and 74.82% of NIH3T3 ions. Comparable %CVs were observed when grouping data by slide (**Suppl. Figure 2E**). A wide range of %CV was still observed, indicating that high %CVs are not attributable to batch effects.

The presence of ions exhibiting low %CVs consistently for both cell lines (**Figure 3F**) suggests that ion variability may be intrinsic to its detectability in HT SpaceM, for example due to abundance or ionization efficiency. Amino acids exhibited the lowest %CVs, with median values less than 5% for both HeLa and NIH3T3 separately (**Figure 3G**). Overall, metabolite classes show similar variability distribution for ions with %CV≤20% for HeLa and NIH3T3, despite cell-specific changes in ion intensities.

In summary, we repurposed correlation, slope, and coefficient of variation metrics to assess the technical reproducibility of HT SpaceM SCM and elucidate ion variability in this framework, aspects little explored by the field. These metrics provided valuable insights into HT SpaceM data quality, highlighting high reproducibility among wells and slides and low variability achieved for a majority of metabolites from diverse metabolic classes.

### HT SpaceM achieves molecular profile and ion variability comparable to bulk LC-MS/MS

While single-cell data offers insights into cell heterogeneity, it is essential to compare conclusions drawn from SCM with those from bulk metabolomics, particularly LC-MS/MS. Moreover, SCM acquired through MALDI imaging MS requires structural validation, which can be provided by LC-MS/MS. As part of our data analysis framework, we evaluated the biological relevance of the metabolite profiles detected by HT SpaceM by comparing HeLa and NIH3T3 cells. UMAP plots show distinct differences between cells from these two lines with the unsupervised Leiden clustering (**Figure 3H**) outweighing technical slide-slide variability (**Figure 3I**). No batch correction methods were necessary for integrating data from different slides. Through differential analysis, the biological relevance of the detected metabolic profiles was further highlighted by the majority (56%) of detected metabolites exhibiting significant relative intensity changes between HeLa and NIH3T3 (FC≥1.5) (**Figure 3J**). More metabolites were found significantly increased in HeLa compared to NIH3T3. Metabolites with high intensities, such as histidine, N-acetylaspartic acid, taurine, and threonine (**Figure 3K**) had the highest Wilcoxon scores.

After establishing the biological relevance of the single-cell metabolic profiles, we validated them with bulk LC-MS/MS. First, we performed structural validation of the putative metabolite identifications from SCM (**Figure 2G**). Second, we compared the variability of metabolite intensities between HT SpaceM and bulk LC-MS/MS (**Figure 3L**). Although 75% of the metabolites demonstrated CV≤20% for both methods (non-zero cell fraction), we also observed differences. Around 16.3% of the metabolites showed low variability (CV≤20%) for bulk but not for single cells, and 7.6% of the metabolites showed low variability for single cells but not for bulk. Single-cell UMAP only for molecular formulas validated with bulk LC-MS/MS and with CV≤20% (n=59) still shows clear discrimination between HeLa and NIH3T3 lines (**Figure 3M**), thus validating the differences demonstrated by the profiles of all metabolites detected in single cells. Furthermore, we compared the levels of single-cell metabolic markers of HeLa and NIH3T3 (dot plot) with their levels in bulk LC-MS/MS (heat map) (**Figure 3N**). Among the differentially expressed (n=42), most of the molecular formulas (85.7%) showed similar trends between single-cell and bulk levels in at least one ion mode.

In summary, by comparing against bulk LC-MS/MS, we validated chemical and biological properties of the metabolic profiles detected by HT SpaceM and proposed a data analysis framework which can be used in other SCM studies.

### Characterizing cancer cell lines from the NCI-60 drug discovery panel at single-cell level

The NCI-60 drug screening panel of human tumor cell lines has been pivotal in cancer research, extensively characterized by various omics, including bulk metabolomics ^35^. However, no SCM characterization has been performed. To address this and showcase the proposed analyses framework, we applied HT SpaceM to nine cell lines from the panel, alongside HeLa, representing diverse tumor origins, including breast (BT-549 and HS 578T), cervical (HeLa), colon (HT29), lung (NCI-H460 and HOP-62), ovarian (IGR-OV1 and OVCAR-5), renal cancers (A498), and melanoma (MALME-3M). Cell morphology, e.g. size, varied across cell lines (**Suppl. Figure 3A)**. NCI-H460 displayed the smallest median size with the least variability in cell area (346.9 µm^2^, IQR=204.0 µm^2^), while BT-549 exhibited the largest median size and variability (952.6 µm^2^, IQR=592.5 µm^2^) (**Suppl. Figure 3B**). We detected 202 [M-H]^−^ ions co-localized with cells (cf. 135 for HeLa and NIH3T3); 39 to 147 ions per cell were detected (**Figure 4A**), with 74.4% of ions present in over 90% of cells. The cell yield at 42,153 cells in 40 wells (39,575 cells of NCI-60 panel and 2,578 HeLa cells) was comparable to the HeLa/NIH3T3 experiment. The bulk LC-MS/MS data of the NCI-60 panel and HeLa revealed 138 metabolites, corresponding to 120 molecular formulas (**Suppl. Figure 3C**). An overlap of 37% (n=75 molecular formulas) of the SC ions detected by both techniques was observed, comprising 85 metabolites identified at the Level 1 and 5 metabolites annotated at the Level 2 following the notation of the Metabolomics Standards Initiative ^32^ . The following NCI60 subset data analysis was performed considering all detected ions in the single-cell dataset given the metabolite coverage.

**Figure 4.**
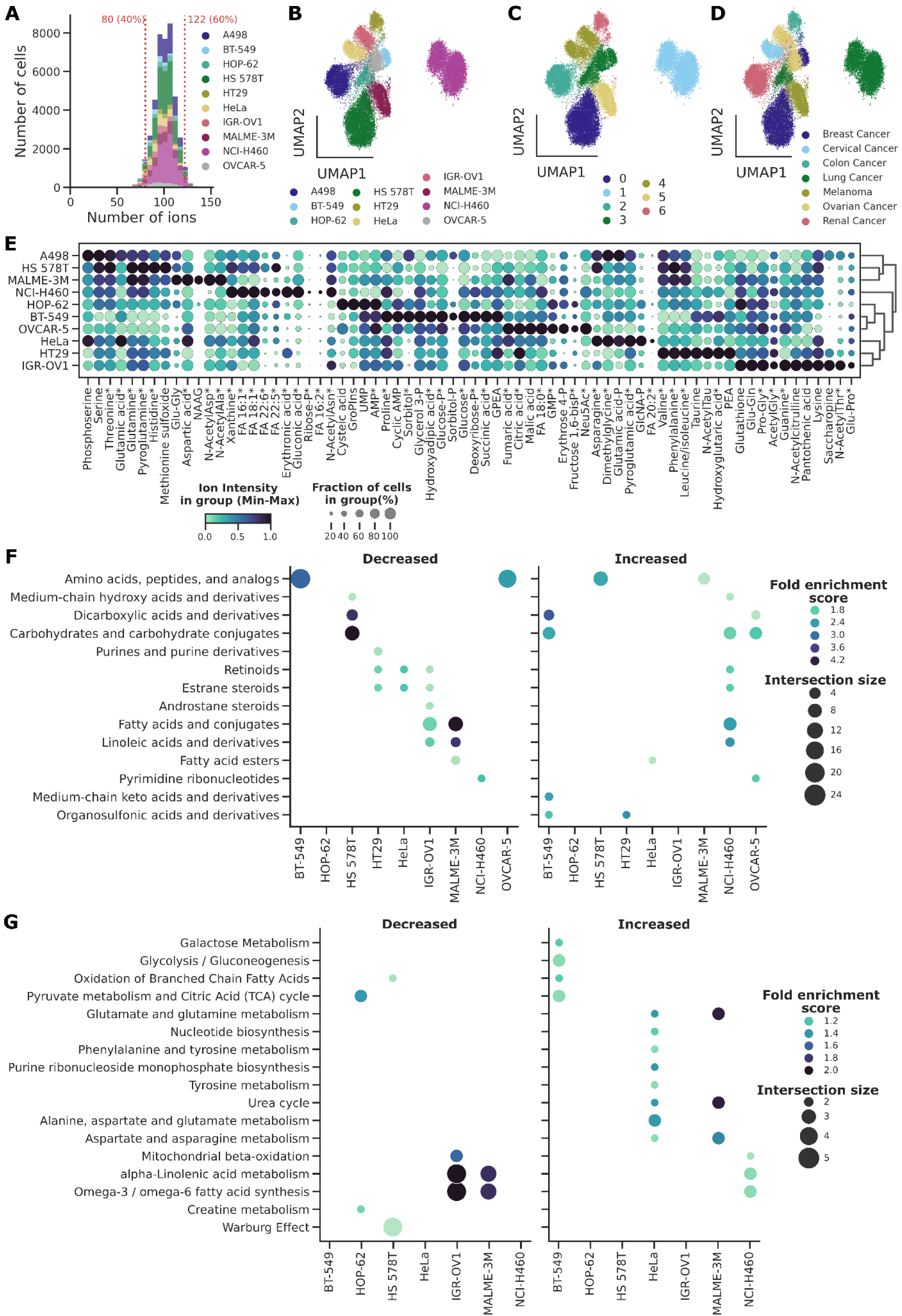
Single-cell characterization of NCI-60 cancer cell lines and HeLa cells from 40 wells. (**A**) Numbers of detected ions per cell, stacked by cell line. (**B**) UMAP of single-cell metabolic profiles from 42,153 cells (39,575 cells from NCI-60 panel and 2,578 HeLa cells) colored by the cell line, considering all detected ions. (**C**) The same as in (**B**) but colored by Leiden clustering. (**D**) The same as in (**B**) but colored by tumor origin. (**E**) Dot plot of cell lines’ markers showing average metabolite intensity feature-scaled and the fraction of cells expressing the metabolite. (**F**) Overrepresentation enrichment analysis of metabolites classes (HMDB) per cell line (*p*-value<0.05). (**G**) Overrepresentation enrichment analysis of metabolic pathways (SPMDB) per cell line (*p*-value<0.1). (**F-G**) Color scale corresponds to the enrichment score and dot size to the number of intersected metabolites with the database; all detected ions in the single-cell analysis were considered for enrichment analysis. Molecule names were validated in bulk LC-MS/MS at Level 1, except as indicated by (*). Metabolites showing isomers in the bulk data are indicated with (**).

UMAP of single-cell metabolic profiles using all detected metabolites shows cells grouping by their cell line (**Figure 4B**), with high reproducibility between the well-replicates (**Suppl. Figure 3D**). Unsupervised Leiden clustering shows clusters comprising more than one cell line (**Figure 4C**), such as the grouping of HT29 and IGR-OV1 in cluster 4, and the grouping of BT-549, OVCAR-5 and HOP-62 in cluster 3. Tumor origin did not influence cell line clustering in the UMAP space (**Figure 4D**). NCI-H460 appears to be distinctly different from other cell lines, including another lung cancer line HOP-62.

Differential analysis (one versus all) revealed distinct metabolite signatures for each line (**Figure 4E**), with cell sizes not biasing metabolite abundance and the discoverability of line markers. Interestingly, some metabolites with high intensities were present in less than 20% of cells (e.g. ribose-phosphate or FA 16:2 in NCI-H460, FA 20:2 in HeLa) demonstrating the unique capacities of SCM. Their high intensity at the same time underscores the biological relevance of zero-intensity cells captured by HT SpaceM, not attributable to instrumental artifacts due to sensitivity limitations. Among the 70 annotated markers displayed in **Figure 4E**, 29 metabolites were not detected in bulk LC-MS/MS analysis at Levels 1 or 2 as indicated (*) and 10 metabolites presented isomers in bulk data (**).

Overrepresentation analysis (ORA) using differential markers exhibited a high abundance and fraction of cells for fatty acids (**Figure 4F**) and fatty acid metabolism (**Figure 4G**) in NCI-H460 cells. This aligns with the reported upregulation of stearoyl-CoA desaturase leading to increased levels of monounsaturated fatty acids (FA 16:1 and FA 18:1) ^36^. Conversely, MALME-3M and IGR-OV1 displayed lower levels of certain fatty acids (FA 16:1, FA 18:1, FA 22:6, FA 22:5) (**Figure 4E**; **Figure 4F**) and significant downregulation in omega-3 and omega-6 fatty acid synthesis and alpha-linolenic acid metabolism (**Figure 4G**), despite originating from different tumor types. Additionally, HS 578T showed downregulation of carbohydrates and derivatives (**Figure 4E**; **Figure 4F**) with low levels of carbohydrate-related metabolism (**Figure 4G**), possibly associated with the observed abnormal activity of the Warburg effect (**Figure 4G**), as breast cancer cells (such as HS 578T) are known for altered glycolysis-related enzymes and transporters ^37^. In contrast, BT-549 exhibited increased carbohydrates and derivatives (**Figure 4E**) with positive overrepresentation of this class (**Figure 4F**). BT-549 presented a trend towards increased pyruvate and citric acid metabolism, and glycolysis/gluconeogenesis (**Figure 4G**).

In summary, HT SpaceM metabolically characterized NCI-60 panel cell lines at the single cell level, with metabolic markers detected even in a small fraction of cells, with ORA providing insights into differential representation of metabolic classes and pathways.

### HT SpaceM reveals single-cell metabolic networks of co-abundant metabolites

Through capturing cell-cell physiological heterogeneity, SCM may uncover the diversity of metabolic programs manifested within cell subpopulations by identifying co-abundant pairs and groups of metabolites within single cells. While analogous methods in single-cell transcriptomics have revealed transcriptional modules through single-cell co-expression analysis ^38^, such exploration has yet to be demonstrated in SCM, likely due to limited small-molecule metabolites coverage and high dropouts rates. Here in this framework, we present an approach for building single-cell metabolic co-abundance networks, showcased for the NCI-60 cancer cell lines.

First, we identified metabolite pairs co-detected across single cells, presenting a low dropout mismatch ratio (16.5% of ion pairs) in each cell line (**Suppl. Figure 3F**). A small proportion of metabolite pairs were substantially co-detected, ranging from 4.7% to 6.7% across cell lines. For each cell line, we computed Pearson correlation (R) between co-detected metabolite pair intensities (**Suppl. Figure 3E**). Most of the co-detected metabolite pairs across cell lines (84.3 to 97.0%) showed weak linear correlation (R<|0.3|) (**Figure 5A**). Co-abundant metabolites, defined as co-detected and showing positive or negative linearity (R>|0.3|) (**Suppl. Figure 3E**), were minority, with positively co-abundant metabolites more prevalent (2.0%-11.0%) than inversely co-abundant metabolites (0.9%-5.2%). IGR-OV1 and OVCAR-5, two ovarian cancer cell lines, exhibited a higher density of positively co-abundant metabolites compared to other cell lines (**Figure 5A**), suggesting strong underlying housekeeping mechanisms of metabolite co-regulation. HeLa was the only line with no strong inversely co-abundant metabolites (R≤-0.5). Some metabolites consistently appeared co-abundant across cell lines (**Figure 5B**), potentially indicating conserved co-regulation, shared pathways, or other mechanisms.

**Figure 5.**
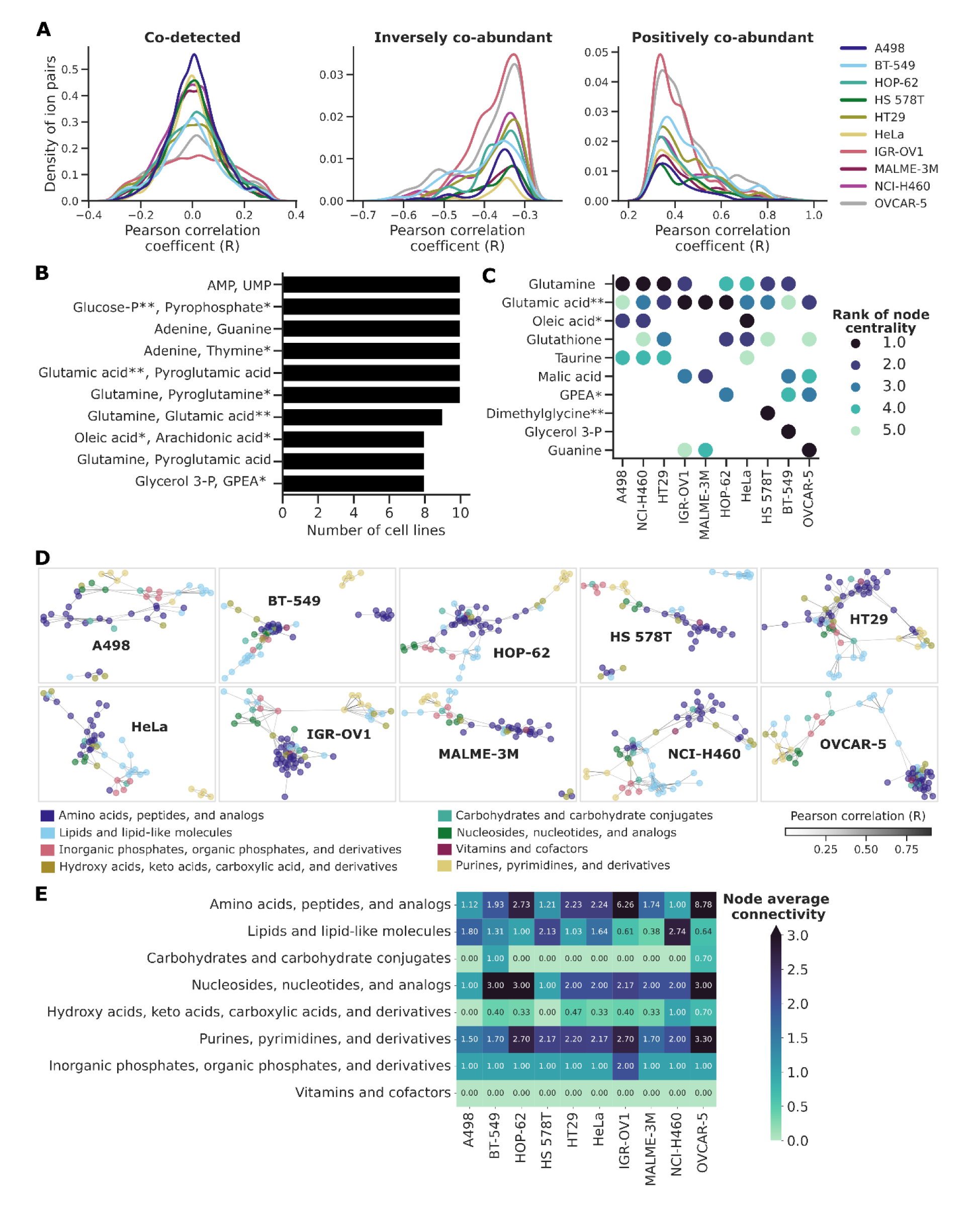
Single-cell metabolite co-abundance networks and their analyses. (**A**) Density distribution of co-detected and co-abundant ion pairs across cell lines, considering all detected ions in the single-cell data. (**B**) Barplot of most frequent (conserved) co-abundant ion pairs across cell lines. (**C**) Dot plot showing metabolite hubs (metabolites top-ranked by their centrality using Pagerank) in the network of each cell line. (**D**) Single-cell metabolite co-abundance networks for each cell line. Node color represents metabolite class and edge represents the Pearson coefficient of correlation (R); only edges with R≥0.3 are shown. (**E**) Heat map of metabolite class-specific node average connectivity per cell line. Molecule names were validated in bulk LC-MS/MS at Level 1 or 2, except as indicated by (*). Metabolites showing isomers in bulk data are indicated with (**)

Next, for each cell line, we built networks connecting (edges) pairs of positively co-abundant metabolites (nodes), highlighting their biochemical relationships. By calculating node centrality using Pagerank with edges weighted by Pearson correlation (**Figure 5C**), we selected the five highest ranked metabolites to identify key ‘hub’ metabolites that were most connected to others in the network for each cell line (**Figure 5D**). Cell lines such as IGR-OV1 and MALME-3M shared similar hub metabolites, including glutamic acid, malic acid, and guanine (**Figure 5C**). Interestingly, these lines also showed similar results in the differential ORA, indicating decreased levels of fatty acids and conjugates, and linoleic acids (**Figure 4E**), along with reduced linoleic acid metabolism and omega fatty acid synthesis (**Figure 4G**). Additionally, A498 and NCI-H460 demonstrated shared metabolite hubs: glutamine, oleic acid, taurine, and glutamic acid. High centrality of glutamine and glutamic acid was observed across various cell lines (**Figure 5C**), with their co-abundance conserved in 9 out of 10 cell lines (**Figure 5B**), suggesting a mechanism for a relatively strict co-regulation of the levels of these enzymatically related molecules. Some metabolic hubs were cell line-specific: BT-549 uniquely presented glycerol 3-phosphate with a high centrality, linking carbohydrate and lipid metabolism in this line. Interestingly, both carbohydrate (glycolysis/gluconeogenesis, citric acid cycle) and lipid (oxidation of branched-chain fatty acids) metabolic pathways were overrepresented in BT-549 (**Figure 4G**).

Expanding our network analysis to metabolite classes, we quantified biochemical relationships between metabolites within each class by calculating class-specific subnetwork node average connectivity (**Figure 5E**). Surprisingly, substantial differences emerged between cell lines. NCI-H460 exhibited a high lipid connectivity (**Figure 5E**), aligning with the upregulation of fatty acids (**Figure 4E**, **Figure 4F**) and fatty acid metabolism (**Figure 4G**), and suggesting single-cell metabolic co-regulation. Similarly, BT-549 showed the highest average connectivity among carbohydrates metabolites (**Figure 5E**), mirroring the upregulation of carbohydrates (**Figure 4E**, **figure 4F**) and carbohydrate metabolism (**Figure 4G**). The ovarian cancer cell lines IGR-OV1 and OVCAR-5 showed similar average connectivity for carbohydrates, lipids, and purines/pyrimidines (**Figure 5E**).

These findings, aligned with the differential ORA, justify the approach of building metabolite co-abundance networks, which can serve as standalone tools for uncovering crucial metabolic hubs or co-regulated metabolic programs.

## Discussion

In this work, we have introduced two major advances which enabled us to achieve higher throughput compared to the original SpaceM method ^20^.

First, we redesigned the experimental setup (**Figure 1A**) by using laser-etched slides with removable well chambers. This modification expanded the well count from 8 to up to 40, more efficiently using the seeded cells, allowing for more technical replicates, and providing experimental flexibility. This versatile setup extends beyond single-cell analysis and can be applied to diverse biological samples like imaging of organoids and spheroids. Second, we restructured the SpaceM processing method (**Figure 1D**) to implement several key improvements. Microscopy acquisition of the entire slide and storage of data in the multi-modal omics spatial data format SpatialData ^29^ simplified region-of-interest (ROI) selection, and contextualization through fluorescent markers, multiplexing, and microscopy-based quality control. Streamlined metabolite annotation with METASPACE ^25^ has accelerated data processing and furthermore allowed us to use it for quality control, data visualization and sharing. Using fine-tuned Cellpose ^28^ trained models have improved the cell segmentation, adapting to variations in cell sizes and morphologies within a single experiment. Finally, batch processing of all wells helped minimize user input, reducing interaction with the interface. Overall, these improvements make the method more efficient experimentally and analytically, saving time, and enhancing control over experimental planning.

The second major advance is the detection of small-molecule metabolites. We demonstrated the coverage of a broad range of metabolite classes (**Figure 2G, Suppl. Figure 1C**) comparable to spatial metabolomics using MALDI-imaging MS ^31^. Even though we demonstrated HT SpaceM for the detection of small-molecule metabolites, the method can also be applied for the detection of lipids by changing the matrix and MALDI-imaging acquisition parameters as shown in SpaceM ^20^. Moreover, the metabolite coverage, reproducibility, and markers revealed by HT SpaceM are comparable to conventional analytical methods such as bulk LC-MS/MS, yet providing single-cell information and detecting markers of small subpopulations as low as 20% of cells (**Figures 2, 3**). Since MALDI-imaging used in HT SpaceM does not resolve isomers and isobars as LC-MS/MS, resulting in Level 2 annotation ambiguity, we proposed and demonstrated a comprehensive approach for LC-MS/MS validation. We showed comparable ion variability to bulk LC-MS/MS, with several metabolites exhibiting %CV≤20% (**Figure 3L**). Low ion variability is essential for bioanalytical method validation, as required by FDA guidelines, and helps evaluate HT SpaceM performance relative to emerging single-cell technologies. HT SpaceM requires substantially low amounts of biological material – only about 2,500 cells seeded per replicate – representing at least a 400-fold reduction compared to typical LC-MS/MS samples ^39^. Overall, HT SpaceM provides reliable SCM data, demonstrating its potential and relevance for studies with limited patient material, such as circulating tumor cells, immune cells, or cells obtained from tumor biopsies.

With these HT SpaceM advancements come certain limitations. The stringent cell culture requirements, including the small surface area of each well and limited cell culture media volume, may challenge cellular adhesion, which is crucial for the method’s success. This issue can be partially circumvented by coating slides with adhesion-promoting compounds like poly-L-lysine or fibronectin. Additionally, mounted chambered wells may leak over extended incubation periods, making them less suitable for experiments requiring long cell culture or in-well treatments. While alternative well-chambered systems could accommodate such treatments, they may reduce the method throughput. HT SpaceM also faces limitations in small-molecule metabolites detection when associated with fixation techniques like paraformaldehyde, which can cause metabolite leakage and negatively impact data quality. The increased sample throughput also demands higher precision in sample preparation; for instance, the presence of crystals from the wash buffer can compromise more samples than in lower throughput methods, as smaller well sizes limit the ability to address issues through ROI selection. Nonetheless, the ability to process more wells per slide helps mitigate this limitation, leading to an experiment less susceptible to batch effects, as evidenced by the high reproducibility observed in this study (**Figure 3C, 3D**). By increasing the number of replicates analyzed in a single experiment, HT SpaceM achieves the highest sample throughput among the SCM techniques with the highest cell yield ^6,40,41^. The proposed data analysis framework, aimed at robustly evaluating the quality of SCM data, benefits from increased sample throughput, aligning with the critical need for data reproducibility.

Future improvements of HT SpaceM could focus on developing a more leakage-resistant experimental setup to streamline cell culture management. Adapting HT SpaceM to higher laser resolutions in mass spectrometers could enhance its spatial resolution and broaden its applicability. The significant data sparsity continues to be a challenge when evaluating performance metrics and establishing correlations across ions. Further incorporation of single-cell genomic tools into HT SpaceM data analysis could enhance data quality and interpretability^42^. Implementing these advancements and establishing data analysis frameworks could propel SCM forward, making it more accessible to clinical translational settings and fostering deeper insights into metabolism.

## Methods

### Cell culture

We cultured 11 cell lines for SCM and bulk LC-MS/MS experiments. These included HeLa Kyoto H2B-mCherry, NIH3T3-GFP, and nine cancer cell lines from the NCI60 panel: A498, BT-549, HOP-64, HT29, HS 578T, IGR-OV1, MALME-3M, NCI-H460, and OVCAR-5. All cell lines were grown separately in 10 cm tissue culture dishes using high glucose DMEM media (1x Pen/Strep) from Gibco/ThermoFisher Scientific, supplemented with 10% fetal bovine serum (FBS) (ThermoFisher), 100 U/mL penicillin, 100 μg/ml streptomycin, and 1 mM sodium pyruvate (all from Gibco). The cells were maintained at 37°C with 5% CO_2_. Cells were split at a 1:10 ratio twice a week using 0.25% trypsin-EDTA (Gibco). Cell counts were determined using Trypan Blue staining and counted with the TC10/TC20 Cell Counter from Bio-Rad Laboratories.

### Laser-etched glass slides

We customized Epredia™ SuperFrost Plus™ glass slides by creating a 40-well layout, arranged in 4 columns and 10 rows, with wells measuring 3 mm x 3 mm. A Zeiss MicroBeam microscope (Carl Zeiss AG) was used to laser-etch on the top surface of each slide with a 10X objective, based on a layout of square elements and well labels designed with PalmRobo software (**Figure 1A**). Following the etching process, the slides were cleaned with isopropanol and air-dried completely before use.

### Cell preparation for SCM experiment

Approximately 2,500 cells suspended in 50 µL of cell media were plated into 40 wells of a 64-well proplate (Grace BioLabs) mounted onto the customized Epredia™ SuperFrost Plus™ etched glass slide (**Figure 1B**). The wells containing different cell lines were randomized across all experiments, with three slides comprising only HeLa and NIH3T3 cells and one slide containing the NCI60 panel alongside HeLa. After 24 h of incubation to allow for cell adhesion, the media was removed and cells were stained with 4’,6-diamidino-2-phenylindole (DAPI) at a concentration of 1 µg/mL (ThermoFisher Scientific) in Phosphate-Buffered Saline (PBS) for 20 min at room temperature. The slides were then detached from the proplates and washed by quickly dipping them three times in 100 mM ammonium acetate. Residual solvent was evaporated, and the cells were desiccated under vacuum (-0.08 kPa) for 30 min at room temperature.

### Brightfield and fluorescence microscopy of cells

Brightfield and fluorescence images (460 nm for DAPI) of the cells were acquired using a Nikon Ti-E inverted microscope (Nikon Instruments) equipped with a Plan Fluor 10X objective (NA 0.30, Nikon Instruments). The images were captured with a pixel size of 0.64 μm and a 20% overlap between consecutive tiles to cover the entire slide using the JOB functionality of the Nikon NIS Elements software. The same imaging configurations were applied for pre- and post-MALDI images (**Figure 1C**).

### Image processing and analysis

Microscopy tiles were stitched into a complete image using the Big Stitcher plug-in ^43^ (**Figure 1D**). The resulting images were stored in SpatialData format ^29^. For selecting MALDI acquisition areas, we used a custom interface written in napari ^44^.

For cell segmentation in the pre-MALDI images, we employed Cellpose ^28^, a deep learning-based method. The *cyto2* model was fine-tuned to accommodate variations in cell size and morphology on a small number of cells manually segmented based on their overlay of brightfield and DAPI images.

For the post-MALDI images, ablation marks were detected and segmented using manual grid fitting with circular shapes used as approximations of the ablation marks.

### MALDI-imaging MS

Before MALDI acquisition, the slides containing cells were sprayed with 1,5-diaminonaphthalene (DAN) (Sigma-Aldrich) prepared at a concentration of 7 mg/mL in 70% acetonitrile (v/v). Spraying was performed in an HTX TM sprayer (HTX Technologies LLC) with the following parameters: temperature at 80°C, 8 passes, a flow rate of 0.05 mL/min, velocity of 1,350 mm/min, track spacing of 3 mm, CC pattern, pressure of 10 psi, gas flow rate of 5 L/min, drying time of 15 seconds, and nozzle height of 41 mm. The estimated matrix density was 0.00069 mg/mm².

For MALDI acquisition, we used a Q-Exactive Plus mass spectrometer (ThermoFisher Scientific) equipped with an AP-SMALDI5 source (Transmit). For the HeLa/NIH3T3 experiment we analyzed one full slide (40 wells) and 2 partial slides (16 wells each), keeping the 1:1 proportion of the number of wells per cell line. For the NCI-60 panel with HeLa, we analyzed a full slide (40 wells) containing 4 wells for each cell line. The analysis was performed in negative ion mode with a mass range of *m/z* 100–400 and a resolving power of 140,000 at *m/z* 200. Acquisition areas of 80 x 80 were interactively selected in each well to target the best regions containing cells. The step size was 25 μm, and the MALDI laser attenuator was set to 29°. For HeLa and NIH3T3 experiment MS parameters included an S-lens voltage of 50 eV, a capillary temperature of 250°C, and a spray voltage of 3.25 kV. For the NCI-60 panel and HeLa experiment MS parameters included a S-lens voltage of 70 eV, a capillary temperature of 350°C, and a spray voltage of 3.1 kV.

### Data pre-processing and metabolite annotation

MALDI-imaging acquisition files were split into individual wells, converted to final centroided imzML files using imzMLConverter ^45^, and uploaded to METASPACE in batch mode (**Figure 1D**). METASPACE, a cloud software (https://metaspace2020.eu), performed metabolite annotation using false discovery rate-controlled annotation ^25^ with an *m/z* accuracy of 3 ppm. Ions were annotated against HMDB (Human Metabolome Database, v4) ^26^ and CoreMetabolome v3 ^27^ databases. Custom databases of molecular formulas were created by selecting ions co-localized with cells by using the area-Normalized Manders Colocalization Coefficient, NMCC, and datasets were reannotated on METASPACE.

### Single-cell data processing

Conversion of MALDI-imaging pixel data into single-cell data, the so-called single-cell data processing, was done as for SpaceM, as described earlier ^20^. After all input data was generated (**Figure 3D**), i.e. pre- and post-MALDI stitched microscopy images, and ion images annotated on METASPACE, the HT SpaceM processing was performed in batch mode to handle the data efficiently. Pre- and post-MALDI images were registered using laser-etched fiducials for improved accuracy and automatization. Segmented cells were aligned to grid-fitted ablation marks, which allowed us to deconvolve pixel-ion intensities provided by METASPACE into single-cell ion intensities. The deconvolution process used the biggest overlap, considering only ablation marks that have at least 30% overlap with cells.

### LC-MS/MS sample preparation

Each cell line was cultured in tissue culture dishes until reaching confluence, yielding approximately 1.5 million cells per replicate (n=3 or 5). After aspirating the cell culture media, the cells were washed by pouring PBS three times over the dish. Once the PBS was discarded, the dish was placed on ice, and 1 mL of ice-cold 80% (v/v) methanol containing isotope-stable internal standards (0.5% final concentration) was added. The plate was then incubated at -80°C for 20 min. Following the incubation, cells were scraped and transferred, along with the solvent mixture, into small tubes. The cell lysate was vortexed at maximum speed for 5 minutes. Further homogenization was performed on dry ice using a bead beater (FastPrep-24; MP Biomedicals, CA, USA) at 6.0 m/s, with three 30-second bursts and a 5-minute pause in between, using 1.0 mm zirconia/glass beads (Biospec Products, OK, USA). After homogenization, the samples were centrifuged for 10 min at 15,000 × g and 4 °C. The supernatants were transferred and dried under a stream of nitrogen (Organomation Microvap, MA, USA). The dried samples were then reconstituted in 100 µL of a solution containing acetonitrile, methanol, and water (2:2:1, v/v), vortexed, and centrifuged under the same conditions. Finally, the supernatants were transferred to silanized glass vials and injected into the analytical system.

### Bulk LC-MS/MS analysis

LC-MS/MS analysis was performed according to the EMBL-MCF 2.0 method ^46^. We used a Vanquish UHPLC system coupled with an Orbitrap Exploris 240 high-resolution mass spectrometer (Thermo Fisher Scientific, MA, USA), operating in both negative and positive electrospray ionization (ESI) modes. Chromatographic separation was achieved on an Atlantis Premier BEH Z-HILIC column (Waters, MA, USA; 2.1 mm × 100 mm, 1.7 µm) at a flow rate of 0.25 mL/min. The mobile phases comprised water (9:1, v/v) for phase A and acetonitrile (9:1, v/v) for phase B, each modified with 10 mM ammonium acetate for negative mode and 10 mM ammonium formate for positive mode. The aqueous components of the mobile phases were pH-adjusted to pH 9.0 (negative mode) with ammonium hydroxide and pH 3.0 (positive mode) with formic acid. The gradient used for separation, including re-equilibration, was as follows: 0 min at 95% B, 2 min at 95% B, 14.5 min at 60% B, 16 min at 60% B, 16.5 min at 95% B, and 20 min at 95% B. The column temperature was maintained at 40°C, the autosampler at 4°C, and the sample injection volume was 5 µL. For MS analysis, full scans were conducted with a mass resolving power of 120,000 over a range of 60–900 *m/z*, with a scan time of 100 ms and an RF lens setting of 70%. MS/MS fragment spectra were obtained through data-dependent acquisition, with a resolving power of 15,000, a scan time of 22 ms, stepped collision energies of 30%, 50%, and 70%, and a cycle time of 900 ms. Ion source parameters included a spray voltage of 4100 V (positive mode) or -3500 V (negative mode), sheath gas at 30 psi, auxiliary gas at 5 psi, sweep gas at 0 psi, ion transfer tube temperature at 350°C, and vaporizer temperature at 300°C.

Samples were measured in a randomized order. Pooled quality control (QC) samples were prepared by combining equal aliquots from each processed sample type. Multiple QC samples were injected at the beginning of the analysis to equilibrate the analytical system, and a QC sample was analyzed after every fifth experimental sample to monitor instrument performance throughout the sequence. To account for background signals, an additional processed blank sample was recorded. Data processing was performed using MS-DIAL 4.9 ^47^, and raw peak intensity data was exported for the annotated molecules and normalized by total ion count for relative metabolite quantification ^48^. Intensities were further log-transformed (log(X+1)).

### Single-cell data analysis framework

The spatio-molecular matrices generated from HT SpaceM processing were used for single-cell data analysis (**Figure 1E**). We filtered the data to include only deprotonated ions [M-H]^−^, retaining cells with at least 20 ions and ions in at least 50 cells. The filtering resulted in a final matrix of 135 ions x 78,500 cells for the HeLa/NIH3T3 dataset and 202 ions × 42,153 cells for the dataset composed of NCI-60 panel cell lines plus HeLa. Ion intensities were normalized to the total ion count per cell, excluding highly expressed ions accounting for more than 5% of the total ion count in a cell. Intensities were scaled to 10,000 during normalization and transformed using the natural logarithm (log(X+1)) before analysis.

### Unsupervised characterization

Principal Component Analysis (PCA) was performed for dimensionality reduction. Using the top 50 PCs, we build a nearest neighbors graph (n_neighbors=15, metric=‘euclidean’) and visualize it through UMAPs colored by cell line, Leiden clustering (unsupervised), replicates and/or tumor origin. No batch correction method was applied to integrate samples from different slides. These processing steps and analyses were conducted using the Scanpy v1.9.1 package ^49^.

Annotated ions were matched to HMDB (www.hmdb.ca) and Kegg (Kyoto Encyclopedia of Genes and Genomes) (www.genome.jp/kegg/) identifiers to evaluate metabolite classes and pathways.

### Differential analysis and enrichment analysis

Single-cell differential analysis considering all annotated metabolites was performed using the Wilcoxon test, with a significant *p*-value<0.05 (FDR-adjusted, two-tailed). Metabolite enrichment analysis was conducted with the R package S2IsoMEr ^50^. We used the simple ORA function considering differentially expressed markers (Log_2_FC>1.5 and *p*-value<0.05) for one cell line versus all. We set the custom_universe as all molecules detected in our analysis, with min_intersection=2 and alpha_cutoff=1 for the matching between the selected markers and the metabolites and metabolic pathways of SMPDB background (Small Molecule Pathway Database (www.smpdb.ca).

### Structural validation

Bulk LC-MS/MS data was annotated against our recently published metabolomics library (EMBL-MCF 2.0) ^46^. Level 1 feature identification used accurate mass, isotope pattern, MS/MS fragmentation, and retention time information, with a minimum matching score of 80%. Metabolites were considered as Level 2 annotations when MS/MS matching scores were less than 80%.

A metabolite was structurally validated when its SCM molecular formula matched with molecular formulas detected by bulk LC-MS/MS in the negative or positive ionization modes. In cases where two or more different metabolites were identified in bulk analysis for one molecular formula, single-cell intensity values for that molecular formula were replicated. The ion intensity of a molecule detected at Level 1 and/or in the negative mode was preferably used for comparison to single-cell metabolite intensities.

### Well-to-well and slide-to-slide reproducibility, and metabolite variability

Reproducibility across wells was evaluated through their correlation. First, we averaged the intensities of each ion across cells within a well replicate, considering both zero and non-zero cell fractions. Then, we calculated the Pearson correlation coefficient (R) between wells of the same condition within the same slide (intra-slide) and between wells from different slides (inter-slide). We also calculated the slope of the best fitting line for the data between two wells in the same manner. For slide-to-slide correlations, we averaged the ion intensities of wells from the same slide for each condition before computing R across slides.

Single-cell ion variability was assessed using the coefficient of variation (%CV) metric. We first averaged the intensities of each ion across cells within a replicate, considering only cells with non-zero intensity values. After obtaining the mean intensity per ion per replicate, we calculated the inter-replicate mean (M) and standard deviation (SD). For bulk data, we calculated M by averaging the intensities of sample replicates within the cell line and computed the respective SD. Metabolite %CV within cell lines was calculated as (SD/M) × 100.

In cases where two or more different metabolites were identified in bulk analysis for one molecular formula, %CV values of single-cell ions were replicated.

### Metabolites correlation and network analysis

Using the single-cell data, we determined within each cell line which metabolites were found co-detected, i.e. metabolites simultaneously detected in a single cell with a low mismatch dropout rate. We selected ion pairs with less than 5% of the cells having zero intensity values simultaneously and more than 60% of cells having non-zero values simultaneously. For those co-detected ions, we computed the Pearson correlation coefficient (R) across metabolite pairs within a cell line, observing if they were positively co-abundant (R>0) or inversely co-abundant (R<0). Network analysis and associated metrics were generated in Python using functions of NetworkX v2.8.6 ^51^. We retained metabolite pairs with R>0.5 to generate network graphs for each cell line, where metabolites (nodes) are colored by metabolite class and a connection between two nodes (edges) represent R. We computed node centrality using the Pagerank algorithm considering R as weight for the edges and calculated the average connectivity within nodes of the same metabolite class for each cell line graph. .

### Statistics and data visualization

For statistical analyses and metrics, we used the Python packages pandas v1.4.3, numpy v1.26.4, scipy v.1.12.0, and sklearn v.0.0.post1. Pathway integrated visualization was performed using iPath3.0 ^33^. For data visualization, we used the Python packages Scanpy v1.9.1, Seaborn v0.13.2, and Matplotlib v3.5.3. Cell images were processed using ImageJ 1.53q (FIJI)^52^. Illustrations and multi-panel figures were created with Inkscape v1.2.1 (www.inkscape.org).

## Data Availability

All annotated MALDI-Imaging data is available at METASPACE (https://metaspace2020.eu/project/HTSpaceM). MALDI-Imaging acquisition files, processed single-cell data, and LC-MS/MS data will be available at MetaboLights (www.ebi.ac.uk/metabolights/MTBLS11236, study identifier MTBLS11236).

## Code Availability

The code used for reproducing figures will shortly be available at GitHub (https://github.com/delafior/HT-SpaceM).

## Acknowledgments

We thank Nicola Zamboni and the National Cancer Institute (NCI)’s contribution in their support for providing the NCI-60 Anti-Cancer Cell Line Panel. T.A. acknowledges funding from the European Research Council (Consolidator grant agreement no. 773089 and Proof-of-Concept grant agreement no. 101101077), Swiss National Science Foundation (Sinergia grant PROMETEX), Michael J. Fox Foundation, and German Research Foundation (DFG).

## Author information

### Contributions

Conceptualization, T.A., M.S., J.D., A.E. Methodology, M.S., A.E., V.H., A.B., B.W., M.E., A.M.,

T.A. Investigation, M.S., B.D. Formal analysis, Data curation and Visualization, J.D. Writing - Original draft J.D., M.S., T.A. Writing - Review and editing, critical input from all authors. Supervision - T.A.

## Ethics declarations

### Declaration of interests

T.A. has a patent application on single-cell metabolomics, leads creation of a startup on single-cell metabolomics incubated at the BioInnovation Institute (BII) in Copenhagen, Denmark, and has a consultancy contract with BII. Other authors declare no competing interests.

## Supplementary Figures

**Supplementary Figure 1.**
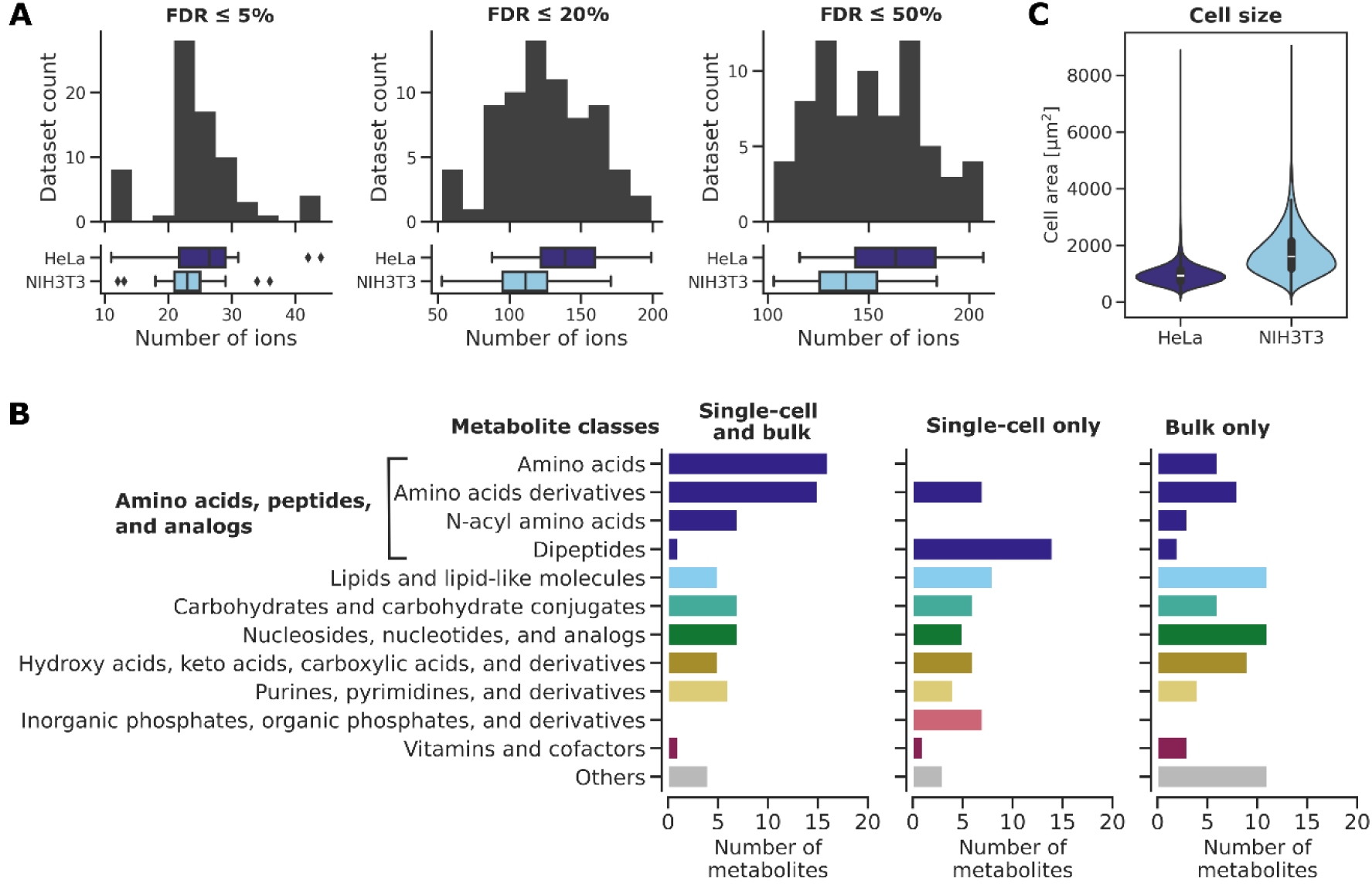
(**A**) Number of ions [M-H]^−^ per well at FDR 5, 20 and 50%. (**B**) Number of metabolites per metabolite class detected in both bulk LC-MS/MS and HT SpaceM, only in single-cell data and only in bulk data. (**C**) Distribution of cell size per cell line.

**Supplementary Figure 2.**
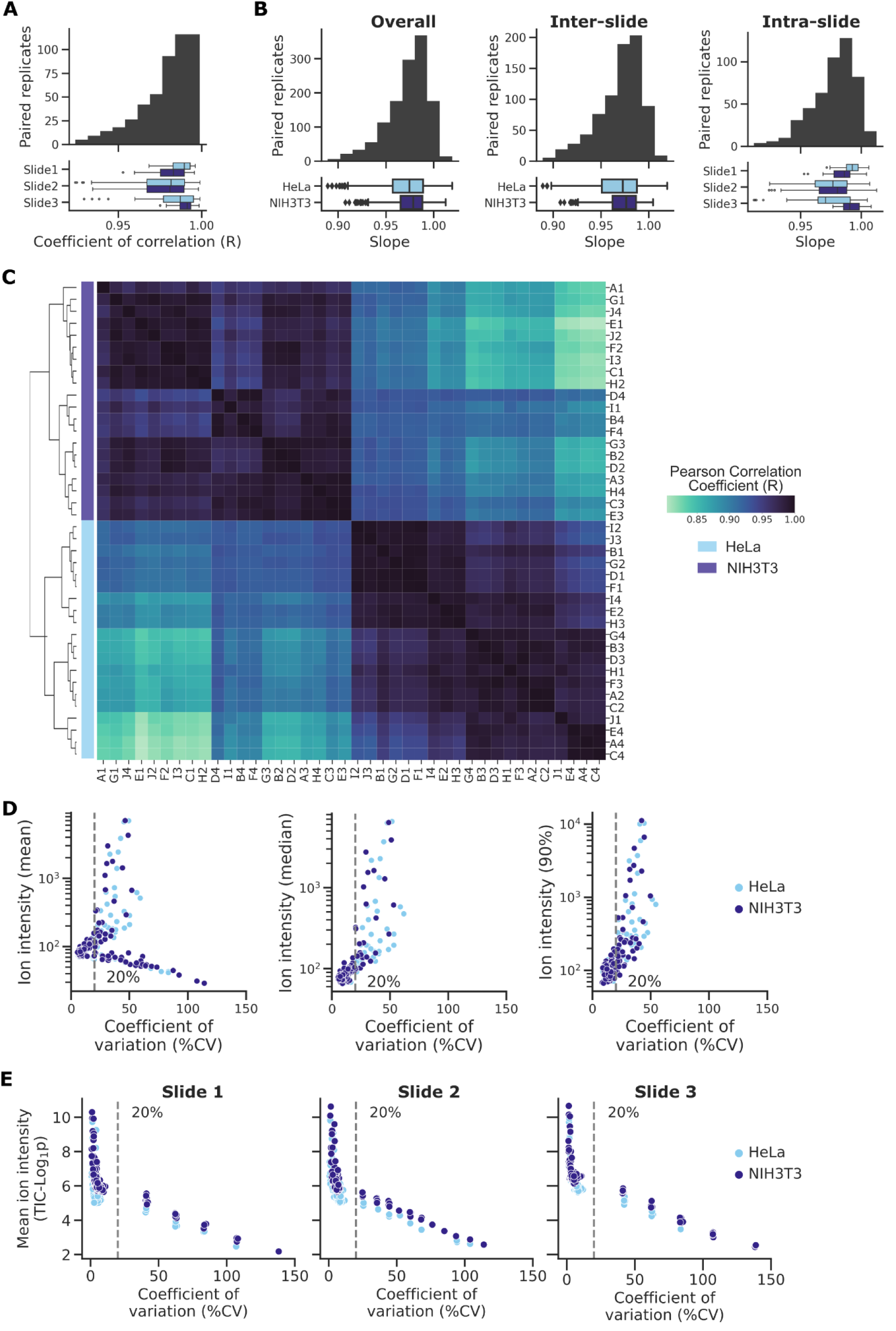
(**A**) Pearson Coefficient of Determination (R^2^) for intra-slide replicates, considering slides and lines separately. (**B**) Slopes distribution for overall, inter-slide and intra-slide replicates. (**C**) Pearson Correlation Coefficient (R) among wells of slide 2. (**D**) Mean, median and 90% quartile ion intensity over Coefficient of Variation (%CV) for HeLa and NIH3T3. (**E**) Normalized mean ion intensity over %CV for HeLa and NIH3T3, per slide.

**Supplementary Figure 3.**
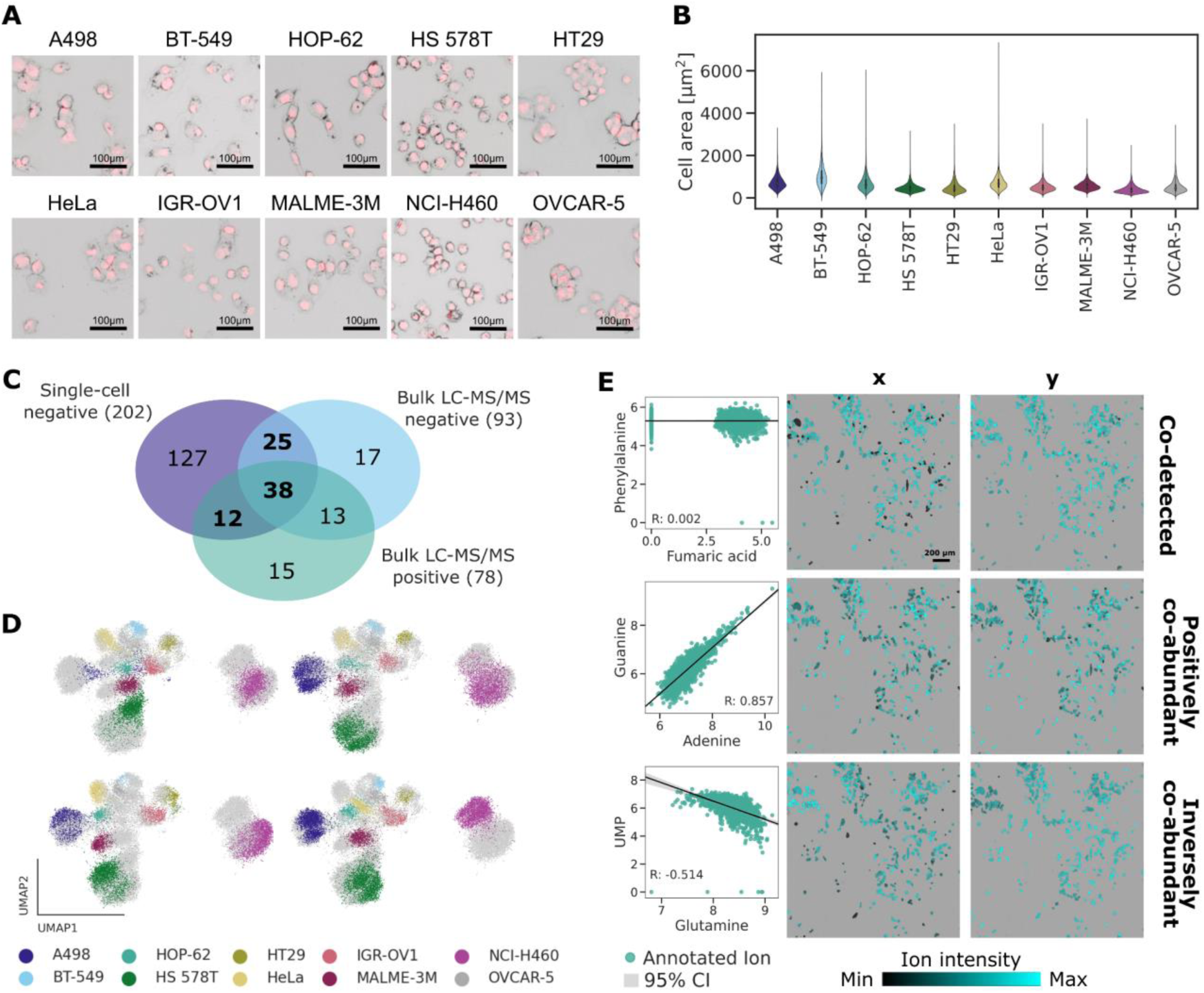
(**A**) Brightfield and fluorescence (DAPI, in red) microscopy images of the NCI60 cells set, alongside HeLa cells. (**B**) Distribution of cell size per cell line. (**C**) Venn diagram highlighting the total number of ions detected by HT SpaceM (negative ion mode) and bulk LC-MS/MS (negative and positive ion modes). (**D**) UMAP of the single-cell data colored by cell line for each replicate (rows, columns). (**E**) Examples of co-detected, positively co-abundant and inversely co-abundant metabolites. Scatter plot of single-cells TIC-normalized ion intensities (Log_1_p) between two metabolites. Microscopy images highlighting the metabolite intensity per cell (x- and y-axis of the scatter plot).

## References

1. Baysoy, A., Bai, Z., Satija, R., and Fan, R. (2023). The technological landscape and applications of single-cell multi-omics. Nat. Rev. Mol. Cell Biol. 24, 695–713.

2. Van de Sande, B., Lee, J.S., Mutasa-Gottgens, E., Naughton, B., Bacon, W., Manning, J., Wang, Y., Pollard, J., Mendez, M., Hill, J., et al. (2023). Applications of single-cell RNA sequencing in drug discovery and development. Nat. Rev. Drug Discov. 22, 496–520.

3. Tirosh, I., and Suva, M.L. (2024). Cancer cell states: Lessons from ten years of single-cell RNA-sequencing of human tumors. Cancer Cell 42, 1497–1506.

4. Zenobi, R. (2013). Single-cell metabolomics: analytical and biological perspectives. Science 342, 1243259.

5. Rubakhin, S.S., Lanni, E.J., and Sweedler, J.V. (2013). Progress toward single cell metabolomics. Curr. Opin. Biotechnol. 24, 95–104.

6. Zhang, C., Le Dévédec, S.E., Ali, A., and Hankemeier, T. (2023). Single-cell metabolomics by mass spectrometry: ready for primetime? Curr. Opin. Biotechnol. 82, 102963.

7. Lanekoff, I., Sharma, V.V., and Marques, C. (2022). Single-cell metabolomics: where are we and where are we going? Curr. Opin. Biotechnol. 75, 102693.

8. Hu, R., Li, Y., Yang, Y., and Liu, M. (2023). Mass spectrometry-based strategies for single-cell metabolomics. Mass Spectrom. Rev. 42, 67–94.

9. Alexandrov, T. (2020). Spatial Metabolomics and Imaging Mass Spectrometry in the Age of Artificial Intelligence. Annu. Rev. Biomed. Data Sci. 3, 61–87.

10. Ma, X., and Fernández, F.M. (2022). Advances in mass spectrometry imaging for spatial cancer metabolomics. Mass Spectrom. Rev., e21804.

11. Saunders, K.D.G., Lewis, H.-M., Beste, D.J.V., Cexus, O., and Bailey, M.J. (2023). Spatial single cell metabolomics: Current challenges and future developments. Current Opinion in Chemical Biology 75, 102327.

12. Priyadarshani, P., Van Grouw, A., Liversage, A.R., Rui, K., Nikitina, A., Tehrani, K.F., Aggarwal, B., Stice, S.L., Sinha, S., Kemp, M.L., et al. (2024). Investigation of MSC potency metrics via integration of imaging modalities with lipidomic characterization. Cell Rep. 43, 114579.

13. Dong, J., Croslow, S.W., Lane, S.T., Castro, D.C., Blanford, J., Zhou, S., Park, K., Burgess, S., Root, M., Cahoon, E., et al. (2024). Enhancing lipid production in plant cells through high-throughput genome editing and phenotyping via a scalable automated pipeline. Synthetic Biology.

14. Xie, Y.R., Castro, D.C., Rubakhin, S.S., Trinklein, T.J., Sweedler, J.V., and Lam, F. (2024). Multiscale biochemical mapping of the brain through deep-learning-enhanced high-throughput mass spectrometry. Nat. Methods 21, 521–530.

15. Buglakova, E., Ekelöf, M., Schwaiger-Haber, M., Schlicker, L., Molenaar, M.R., Shahraz, M., Stuart, L., Eisenbarth, A., Hilsenstein, V., Patti, G.J., et al. (2024). Spatial single-cell isotope tracing reveals heterogeneity of de novo fatty acid synthesis in cancer. Nat. Metab. 6, 1695–1711.

16. Blanc, L., Grelard, F., Tuck, M., Dartois, V., Peixoto, A., and Desbenoit, N. (2024). In tissue spatial single-cell metabolomics by coupling mass spectrometry imaging and immunofluorescences. bioRxiv, 2024.03.22.586317. 10.1101/2024.03.22.586317.

17. McKinnon, J.C., Balez, R., Young, R.S.E., Brown, M.L., Lum, J.S., Robinson, L., Belov, M.E., Ooi, L., Tortorella, S., Mitchell, T.W., et al. (2024). MALDI-2-enabled oversampling for the mass spectrometry imaging of metabolites at single-cell resolution. J. Am. Soc. Mass Spectrom. 10.1021/jasms.4c00241.

18. Xu, S., Yang, C., Yan, X., and Liu, H. (2022). Towards high throughput and high information coverage: advanced single-cell mass spectrometric techniques. Anal. Bioanal. Chem. 414, 219–233.

19. Neumann, E.K., Ellis, J.F., Triplett, A.E., Rubakhin, S.S., and Sweedler, J.V. (2019). Lipid Analysis of 30 000 Individual Rodent Cerebellar Cells Using High-Resolution Mass Spectrometry. Anal. Chem. 91, 7871–7878.

20. Rappez, L., Stadler, M., Triana, S., Gathungu, R.M., Ovchinnikova, K., Phapale, P., Heikenwalder, M., and Alexandrov, T. (2021). SpaceM reveals metabolic states of single cells. Nat. Methods 18, 799–805.

21. Liu, R., and Yang, Z. (2021). Single cell metabolomics using mass spectrometry: Techniques and data analysis. Anal. Chim. Acta 1143, 124–134.

22. Pan, X., Pan, S., Du, M., Yang, J., Yao, H., Zhang, X., and Zhang, S. (2024). SCMeTA: a pipeline for single-cell metabolic analysis data processing. Bioinformatics 40, btae545.

23. Comi, T.J., Neumann, E.K., Do, T.D., and Sweedler, J.V. (2017). microMS: A Python Platform for Image-Guided Mass Spectrometry Profiling. J. Am. Soc. Mass Spectrom. 10.1007/s13361-017-1704-1.

24. Ibáñez, A.J., Fagerer, S.R., Schmidt, A.M., Urban, P.L., Jefimovs, K., Geiger, P., Dechant, R., Heinemann, M., and Zenobi, R. (2013). Mass spectrometry-based metabolomics of single yeast cells. Proc. Natl. Acad. Sci. U. S. A. 110, 8790–8794.

25. Palmer, A., Phapale, P., Chernyavsky, I., Lavigne, R., Fay, D., Tarasov, A., Kovalev, V., Fuchser, J., Nikolenko, S., Pineau, C., et al. (2017). FDR-controlled metabolite annotation for high-resolution imaging mass spectrometry. Nat. Methods 14, 57–60.

26. Wishart, D.S., Feunang, Y.D., Marcu, A., Guo, A.C., Liang, K., Vázquez-Fresno, R., Sajed, T., Johnson, D., Li, C., Karu, N., et al. (2018). HMDB 4.0: the human metabolome database for 2018. Nucleic Acids Res. 46, D608–D617.

27. Wadie, B., Stuart, L., Rath, C.M., Drotleff, B., Mamedov, S., and Alexandrov, T. (2023). METASPACE-ML: Metabolite annotation for imaging mass spectrometry using machine learning. bioRxiv. 10.1101/2023.05.29.542736.

28. Stringer, C., Wang, T., Michaelos, M., and Pachitariu, M. (2021). Cellpose: a generalist algorithm for cellular segmentation. Nat. Methods 18, 100–106.

29. Marconato, L., Palla, G., Yamauchi, K.A., Virshup, I., Heidari, E., Treis, T., Vierdag, W.-M., Toth, M., Stockhaus, S., Shrestha, R.B., et al. (2024). SpatialData: an open and universal data framework for spatial omics. Nat. Methods. 10.1038/s41592-024-02212-x.

30. Alexandrov, T., Ovchinnikova, K., Palmer, A., Kovalev, V., Tarasov, A., Stuart, L., Nigmetzianov, R., Fay, D., Key METASPACE contributors, Gaudin, M., et al. (2019). METASPACE: A community-populated knowledge base of spatial metabolomes in health and disease. bioRxiv. 10.1101/539478.

31. Saharuka, V., Vieira, L.M., Stuart, L., Ekelöf, M., Molenaar, M.R., Bailoni, A., Ovchinnikova, K., Soltwisch, J., Bausbacher, T., Jakob, D., et al. (2024). Large-scale evaluation of spatial metabolomics protocols and technologies. bioRxiv. 10.1101/2024.01.29.577354.

32. Sumner, L.W., Amberg, A., Barrett, D., Beale, M.H., Beger, R., Daykin, C.A., Fan, T.W.-M., Fiehn, O., Goodacre, R., Griffin, J.L., et al. (2007). Proposed minimum reporting standards for chemical analysis. Metabolomics 3, 211–221.

33. Darzi, Y., Letunic, I., Bork, P., and Yamada, T. (2018). iPath3.0: interactive pathways explorer v3. Nucleic Acids Res. 46, W510–W513.

34. Stegle, O., Teichmann, S.A., and Marioni, J.C. (2015). Computational and analytical challenges in single-cell transcriptomics. Nat. Rev. Genet. 16, 133–145.

35. Cherkaoui, S., Durot, S., Bradley, J., Critchlow, S., Dubuis, S., Masiero, M.M., Wegmann, R., Snijder, B., Othman, A., Bendtsen, C., et al. (2022). A functional analysis of 180 cancer cell lines reveals conserved intrinsic metabolic programs. Mol. Syst. Biol. 18, e11033.

36. Noto, A., Raffa, S., De Vitis, C., Roscilli, G., Malpicci, D., Coluccia, P., Di Napoli, A., Ricci, A., Giovagnoli, M.R., Aurisicchio, L., et al. (2013). Stearoyl-CoA desaturase-1 is a key factor for lung cancer-initiating cells. Cell Death Dis. 4, e947–e947.

37. Shin, E., and Koo, J.S. (2021). Glucose metabolism and glucose transporters in breast cancer. Front. Cell Dev. Biol. 9. 10.3389/fcell.2021.728759.

38. Iacono, G., Massoni-Badosa, R., and Heyn, H. (2019). Single-cell transcriptomics unveils gene regulatory network plasticity. Genome Biol. 20. 10.1186/s13059-019-1713-4.

39. Mackay, G.M., Zheng, L., van den Broek, N.J.F., and Gottlieb, E. (2015). Analysis of cell metabolism using LC-MS and isotope tracers. In Methods in Enzymology Methods in enzymology. (Elsevier), pp. 171–196.

40. Do, T.D., Comi, T.J., Dunham, S.J.B., Rubakhin, S.S., and Sweedler, J.V. (2017). Single cell profiling using ionic liquid matrix-enhanced secondary ion mass spectrometry for neuronal cell type differentiation. Anal. Chem. 89, 3078–3086.

41. Zhang, L., Xu, T., Zhang, J., Wong, S.C.C., Ritchie, M., Hou, H.W., and Wang, Y. (2021). Single cell metabolite detection using inertial microfluidics-assisted ion mobility mass spectrometry. Anal. Chem. 93, 10462–10468.

42. Bilous, M., Hérault, L., Gabriel, A.A., Teleman, M., and Gfeller, D. (2024). Building and analyzing metacells in single-cell genomics data. Mol. Syst. Biol. 20, 744–766.

43. Hörl, D., Rojas Rusak, F., Preusser, F., Tillberg, P., Randel, N., Chhetri, R.K., Cardona, A., Keller, P.J., Harz, H., Leonhardt, H., et al. (2019). BigStitcher: reconstructing high-resolution image datasets of cleared and expanded samples. Nat. Methods 16, 870–874.

44. Sofroniew, N., Lambert, T., Bokota, G., Nunez-Iglesias, J., Sobolewski, P., Sweet, A., Gaifas, L., Evans, K., Burt, A., Doncila Pop, D., et al. (2024). napari: a multi-dimensional image viewer for Python (Zenodo) 10.5281/ZENODO.13863809.

45. Race, A.M., Styles, I.B., and Bunch, J. (2012). Inclusive sharing of mass spectrometry imaging data requires a converter for all. J. Proteomics 75, 5111–5112.

46. Dekina, S., Alexandrov, T., and Drotleff, B. (2024). EMBL-MCF 2.0: an LC-MS/MS method and corresponding library for high-confidence targeted and untargeted metabolomics using low-adsorption HILIC chromatography. Metabolomics 20. 10.1007/s11306-024-02176-1.

47. Tsugawa, H., Cajka, T., Kind, T., Ma, Y., Higgins, B., Ikeda, K., Kanazawa, M., VanderGheynst, J., Fiehn, O., and Arita, M. (2015). MS-DIAL: data-independent MS/MS deconvolution for comprehensive metabolome analysis. Nat. Methods 12, 523–526.

48. Drotleff, B., and Lämmerhofer, M. (2019). Guidelines for selection of internal standard-based normalization strategies in untargeted lipidomic profiling by LC-HR-MS/MS. Anal. Chem. 91, 9836–9843.

49. Wolf, F.A., Angerer, P., and Theis, F.J. (2018). SCANPY: large-scale single-cell gene expression data analysis. Genome Biol. 19. 10.1186/s13059-017-1382-0.

50. Wadie, B., Molenaar, M.R., Vieira, L.M., and Alexandrov, T. (2024). Enrichment analysis for spatial and single-cell metabolomics accounting for molecular ambiguity. bioRxiv. 10.1101/2024.08.23.609355.

51. Hagberg, A., Swart, P.J., and Schult, D.A. (1 2008). Exploring network structure, dynamics, and function using NetworkX. In.

52. Schindelin, J., Arganda-Carreras, I., Frise, E., Kaynig, V., Longair, M., Pietzsch, T., Preibisch, S., Rueden, C., Saalfeld, S., Schmid, B., et al. (2012). Fiji: an open-source platform for biological-image analysis. Nat. Methods 9, 676–682.

